# Asbestos Mineral Fibre Exposure Significantly Affects Volatile Organic Compound Profiles of Mesothelial Cell Lines In Vitro

**DOI:** 10.1101/2024.07.08.602545

**Authors:** Sam Bonsall, Theo Issitt, Nick Peake, Mari Herigstad, Jason Webber, Sarah Haywood-Small

## Abstract

Malignant mesothelioma (MM) is a rare cancer caused by exposure to asbestos, this condition continues to represent a significant diagnostic and therapeutic challenge. Due to the long disease latency, non-invasive diagnostic modalities such as breath analysis may enhance MM early detection by identifying the disease specific pattern of volatile organic compounds (VOCs) in exhaled breath. In this study, solid phase microextraction (SPME) and gas chromatography-mass spectrometry (GC-MS) was used to extract VOCs from the headspace of two mesothelioma cell lines: MSTO-211H (biphasic mesothelioma) and NCI-H28 (epithelioid mesothelioma) in addition to MET-5a (non-malignant mesothelial cell line), following exposure to asbestos mineral fibers (actinolite, amosite, crocidolite and chrysotile) and a non-asbestos fiber control (wollastonite). Multivariate statistical analysis was applied to identify VOC-based biomarkers associated with asbestos exposure within the cell lines. Data confirms that exposure to mineral fibers (including the non-asbestos fiber control) induces significant alterations in the levels of as many as 24 VOCs in each cell line. This is the first study to investigate the impact of asbestos mineral fiber exposure on the VOC profile of MM cell lines. This may be relevant to studies involving mesothelioma patients, as these VOCs could also serve as indicators of asbestos exposure in individuals who have been exposed to asbestos.

## 1. Introduction

Malignant mesothelioma (MM) is a rare, incurable cancer affecting the lining of the lung, primarily associated with occupational asbestos fibre exposure. While asbestos usage is banned in the UK, the mining, importation, and usage of asbestos continues in many countries, despite recent studies directly linking a decreased incidence of MM to asbestos bans [1]

MM represents an ongoing diagnostic challenge. An overwhelming majority of MM cases are diagnosed at an advanced stage of disease progression, precluding patients from receiving curative treatment [2]. The largely non-specific symptoms associated with MM, coupled with the potential 60+ year latency period between initial asbestos exposure and development of malignancy both contribute to the difficulty of detecting MM at an earlier stage. More specific symptoms of MM, such as pleural effusion and shortness of breath arise at a much later stage of disease progression [3]. MM can only be clinically diagnosed via invasive pleural biopsy and thoracentesis at the onset of late-stage symptoms [4]. This highlights the need for a more effective diagnostic method for MM that can detect the disease at an earlier and potentially more treatable stage. Currently, MM has a median survival time between 8- and 14-months following diagnosis [5] Early detection of MM could lead to better patient outcomes and enable further research into potential curative treatments to be conducted on patients with earlier stage disease.

An emerging diagnostic modality for cancer is breath analysis, whereby the levels of endogenous volatile organic compounds (VOCs) are measured in an individual’s breath, usually via gas chromatography/ mass spectrometry (GC/MS). VOCs are low vapor pressure compounds which are released from cells as part of regular metabolic processes. Internal metabolic processes associated with disease can be disturbed, altering the VOC metabolites released and consumed by cells [6,7]. Therefore, individual VOCs or a VOC profile can serve as biomarkers of disease or progression towards a diseased state. In the context of cancer, this is a particularly attractive diagnostic option, as the accumulation of genetic mutations on top of rapid, uncontrolled cellular division may result in distinct VOC profiles related to these characteristics.

While high levels of accuracy and sensitivity have been achieved in studies investigating the validity of prospective diagnostic breath tests for pleural malignancies including MM [8,9], wide scale clinical validation is still required for breath tests to be adopted for use in the diagnosis of cancer. The precise relationship between malignancies and the various volatile organic compounds (VOCs) that have been identified as potential biomarkers is not yet fully understood. In general, cellular stress is thought to result in increased levels of lipid peroxidation, which in turn leads to altered production of VOCs [10,11] which can then be detected via breath test. Furthermore, recent studies have successfully differentiated malignant mesothelial cell lines from non-malignant mesothelial and lung cancer cell lines based on VOC profile alone, indicating that MM induces a VOC profile distinct from both lung cancer and non-malignant mesothelium, which could potentially be detected in an individual’s breath [12,13].

For the first time, this research describes the effect of asbestos mineral fiber (AMF) exposure upon cellular VOC profiles within a controlled, laboratory-based setting. We hypothesize that AMF exposure (actinolite, amosite, crocidolite and chrysotile) and a non-AMF control (wollastonite) will result in significant, detectable changes in VOC profile for mesothelial cell lines (MSTO-211H, biphasic and NCI-H28, epithelioid) and non-malignant mesothelial cell line, MET5a.

## 2. Materials and Methods

### Cell Culture

MET-5a, MSTO-211H and NCI-H28 cells were purchased from ATCC. MSTO-211H and NCI-H28 cells were cultured in RPMI-1640 medium with L-glutamine (Thermo Fisher, Loughborough, UK) supplemented with 10% v/v FBS and 1% v/v Penicillin/Streptomycin (Thermo Fisher). MET-5a cells were cultured in Medium 199 (Merck, London, UK) supplemented with 10% v/v FBS, 1% v/v Penicillin/Streptomycin, 400nM HEPES, 440µM hydrocortisone, 870µM zinc free bovine insulin and 3.3nM human epidermal growth factor as previously described [14]. cell lines were maintained at 37°C in a 5 % CO2 atmosphere. Culture medium was renewed every 3 days. Cells were subcultured at 70-80% confluency by detachment with 0.4% trypsin-EDTA solution. All cell lines were routinely screened throughout the study and found to be negative for mycoplasma with the MycoAlert detection kit (Lonza).

### Headspace gas analysis of mesothelial cell lines

1.8×106 cells were seeded into standard T75 culture flasks and incubated for 24 hours. Culture medium was then discarded and replaced with 15 mL culture medium containing 5µg/mL AMFs. Cells were then incubated in the presence of AMFs for a further 48 hours. A 50/30µM DVB/CAR/PDMS SPME fiber was then used to extract VOCs from culture flask headspace by piercing the cap filter and exposing the SPME fiber to the flask headspace for 15 minutes, then retracting the fiber into the fiber holder assembly. The assembly and filter cap were sealed with parafilm for the duration of sampling. The SPME fiber was cleaned by insertion into a GC-MS inlet for 10 minutes at 250°C prior to VOC extraction. VOC extraction was performed inside a cell culture incubator under standard culture conditions. New SPME fibers were conditioned in a GC-MS inlet at 270°C for 30 minutes in accordance with manufacturer instructions. “Blank” flasks containing only cell culture medium with corresponding AMFs were also analyzed identically.

### Asbestos Mineral Fibres

Asbestos mineral fibers were managed as described previously [15]. The Union for International Cancer Control-accredited fibers (Health and Safety Laboratories, UK, as described in table 1) were prepared in specially designed laminar flow hoods (Santia Asbestos Management Ltd. laboratories) in a phosphate-buffered saline (PBS) solution to a final stock concentration of 2 mg/ml. AMFs pose a minimal risk of inhalation in an aqueous solution and were stored in PBS and regularly checked to ensure they remained in solution. Solutions were sterilized at 121 °C in an autoclave and stored at room temperature, protected from light.

**Table 1.** AMFs used in this study, and their associated morphology.

| Fibre Name | Fibre Morphology |
| --- | --- |
| Actinolite | Amphibole |
| Amosite | Amphibole |
| Crocidolite | Amphibole |
| Chrysotile | Serpentine |
| Wollastonite | Non-Asbestos Fibre Control |

**Table 2.** Significantly altered VOCs (P<0.05 vs untreated) in the MET-5a cell line following AMF exposure.

| MET-5a |  |  |
| --- | --- | --- |
| Average RT | Tentative ID | Functional Group |
| 2.89 | Isopropyl Alcohol(2-D) | Alcohol |
| 2.99 | Isoprene | Alkene |
| 3.73 | Butane | Alkane |
| 7.12 | Isopropenyl Methyl Ether | Ether |
| 8.35 | Benzene | Aromatic |
| 13.81 | Isopentyl Phenylacetate | Aromatic |
| 14.44 | Thymol | Aromatic |
| 16.18 | 2-Hexanone | Ketone |
| 16.97 | Ethylbenzene | Aromatic |
| 17.00 | Benzyl Isovalerate | Aromatic |
| 19.49 | 3-Methyl-1-Pentanol | Alcohol |
| 19.67 | Isopropylbenzene | Aromatic |
| 20.12 | 5-Nonanone | Alkane |
| 20.69 | Undecane | Alkane |
| 20.98 | Propylbenzene | Aromatic |
| 21.34 | Decane | Alkane |
| 21.45 | Hexamethylethane | Alkane |
| 23.11 | 2-Methylheptane | Alkane |
| 23.45 | Tridecane | Alkane |
| 23.88 | Nonacosane | Alkane |
| 24.03 | 3-Phenylindole | Aromatic |
| 24.18 | Octadecane | Alkane |
| 24.25 | Benzaldehyde | Aldehyde |
| 25.06 | Methyl Hexyl Ketone | Ketone |
| 27.85 | 4-Hydroxymandelic Acid | Aromatic |
| 28.08 | Eicosane | Alkane |
| 28.34 | Hexadecane | Alkane |
| 31.79 | Naphthalene | Aromatic |

### GC-MS analysis of VOCs

GC-MS analysis of VOCs was conducted as previously described (Little *et al*., 2020). An Agilent 7890A with an Rtx-VMS capillary column (30 m x 0.25 mm x 1.4 µm; Restek, Saunderton, UK) coupled to a MS-5975C triple axis detector was used for VOC analysis. The GC-MS inlet temperature was set to 250°C for initial injection of the SPME fiber. The oven program was set as follows: Initial temperature of 35°C held for 5 minutes, ramped to 140°C at a rate of 4°C min^-1^ and held for 5 minutes, then finally ramped to 240 °C at a rate of 20°C min^-1^ and held for 4 minutes. This gave a total run time of 45.25 minutes for one sample. The MS transfer line was set to 260°C and VOC analysis was performed in full scan mode with a range of 35-300 a.m.u. VOCs were analyzed via direct injection of the SPME fiber into the GC-MS inlet. The SPME fiber was exposed for 10 minutes after insertion of the fiber assembly needle into the GC-MS inlet to simultaneously release VOCs for analysis and clean the SPME fiber for subsequent VOC extractions.

### Cell Viability

All cells were assayed for viability via trypan blue exclusion test following VOC extraction to confirm that AMF treatments did not negatively impact cell viability. T75 flasks containing cells were briefly washed with PBS, then trypsinized with 0.4% trypsin-EDTA to allow full detachment of cells. Cell viability and count were then assessed using 0.05% trypan blue in a 1:1 ratio with cell suspensions. Cell counts were performed with a countess II automated cell counter (Thermo Fisher).

### Data processing and analysis

Mass spectra were exported from masshunter as CDF files and deconvolved and analyzed using the *eRah* R package [16,17]. Spectra were deconvolved with the parameters: min.peak.width =1,1; avoid.processing.mz = c(35:45), and the resulting deconvolved spectra were then aligned with the parameters: min.spectra.cor = 0.8; max.time.dist = 4; mz.range = 35:300. Metabolites were then identified via spectral match to the MoNA GC/MS spectra database [18]. Significantly different compounds between culture medium and their corresponding cell line/treatment combination were retained, and insignificant compounds were omitted from subsequent analysis. Compounds with less than 80% spectral match were also omitted from subsequent analysis. The resulting aligned data matrix containing tentative compound identities was then analyzed using MetaboanalystR 4.0 [19,20]. Principal component analysis (PCA) and sparse-partial least squares discriminant analysis (s-PLSDA) plots were produced with the default command parameters, and significant compounds associated with each AMF treatment were determined using T-Tests for each individual compound, comparing untreated cells to each AMF exposure group individually.

## 3. Results

### Effect of Asbestos Fibre Stimulation on Cell Viability

Trypan blue exclusion tests were conducted following VOC analysis to ensure asbestos fiber exposure did not affect cell viability. Asbestos fiber stimulation did not significantly affect cell viability or cell number for any cell line or fiber treatment (Figure 1.)

**Figure 1.**
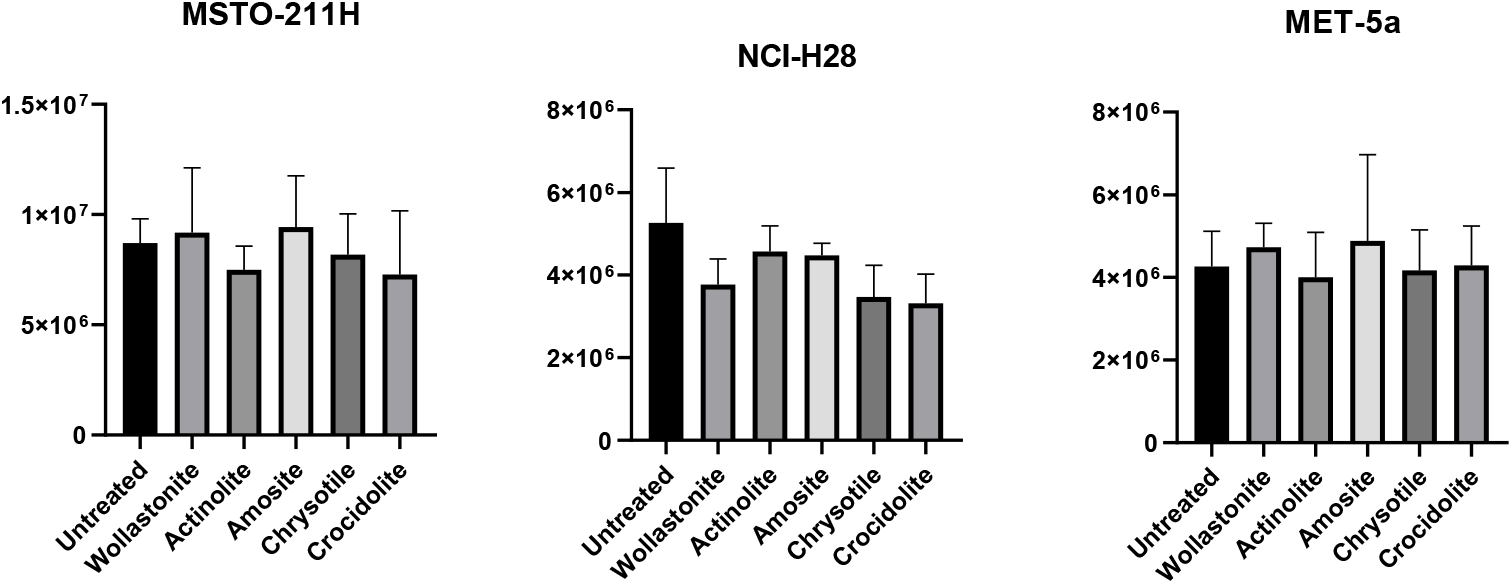
Trypan blue exclusion test total live cell count results for all cell lines stimulated with 5 µg/mL AMF for 48 hours. One way ANOVA analysis did not reveal significant differences in cell number or viability following AMF stimulation (P>0.05).

### PCA and SPLS-DA Analysis

Background VOCs and siloxane compounds originating from cell culture medium, flasks and artefacts from the sampling process were removed prior to data analysis as described previously [12]. Each cell line displayed separation between untreated and AMF treated cells on both PCA and SPLS-DA plots, indicating an alteration in VOC profile is induced following AMF stimulation.

All cell lines showed some degree of separation between AMF treatments following PCA (Figure 2), indicating that AMF exposure induced a different VOC profile compared to untreated cells.

**Figure 2.**
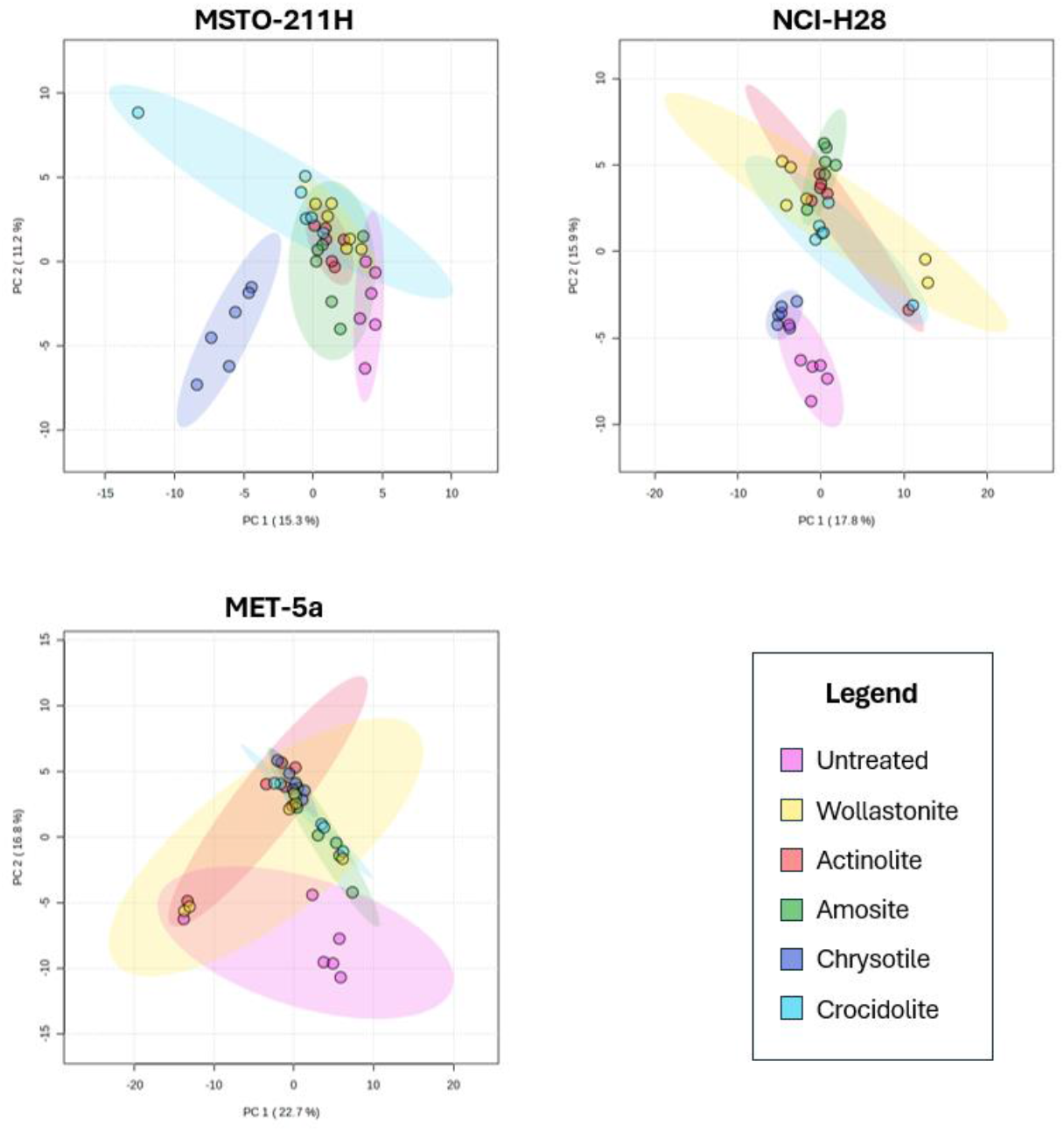
PCA plots of mesothelial cell lines treated with AMFs. Each individual point represents a single experimental repeat (n=6 per treatment).

The MET-5a cell line treatments clustered into two distinct groups, an untreated group and a second, tightly clustered group consisting of every AMF treatment, including the non-asbestos control fiber, wollastonite, suggesting that AMF exposure induces a unique VOC profile in non-malignant mesothelial cells, regardless of the specific mineral fiber used.

The NCI-H28 cell line displayed three main groups. The first consisted of untreated cells, and the second of chrysotile treated cells, with minor overlap of a single experimental repeat from the untreated cells between these two groups. The third group consisted of relatively tightly clustered points from all remaining AMF treatments (Figure 3).

**Figure 3.**
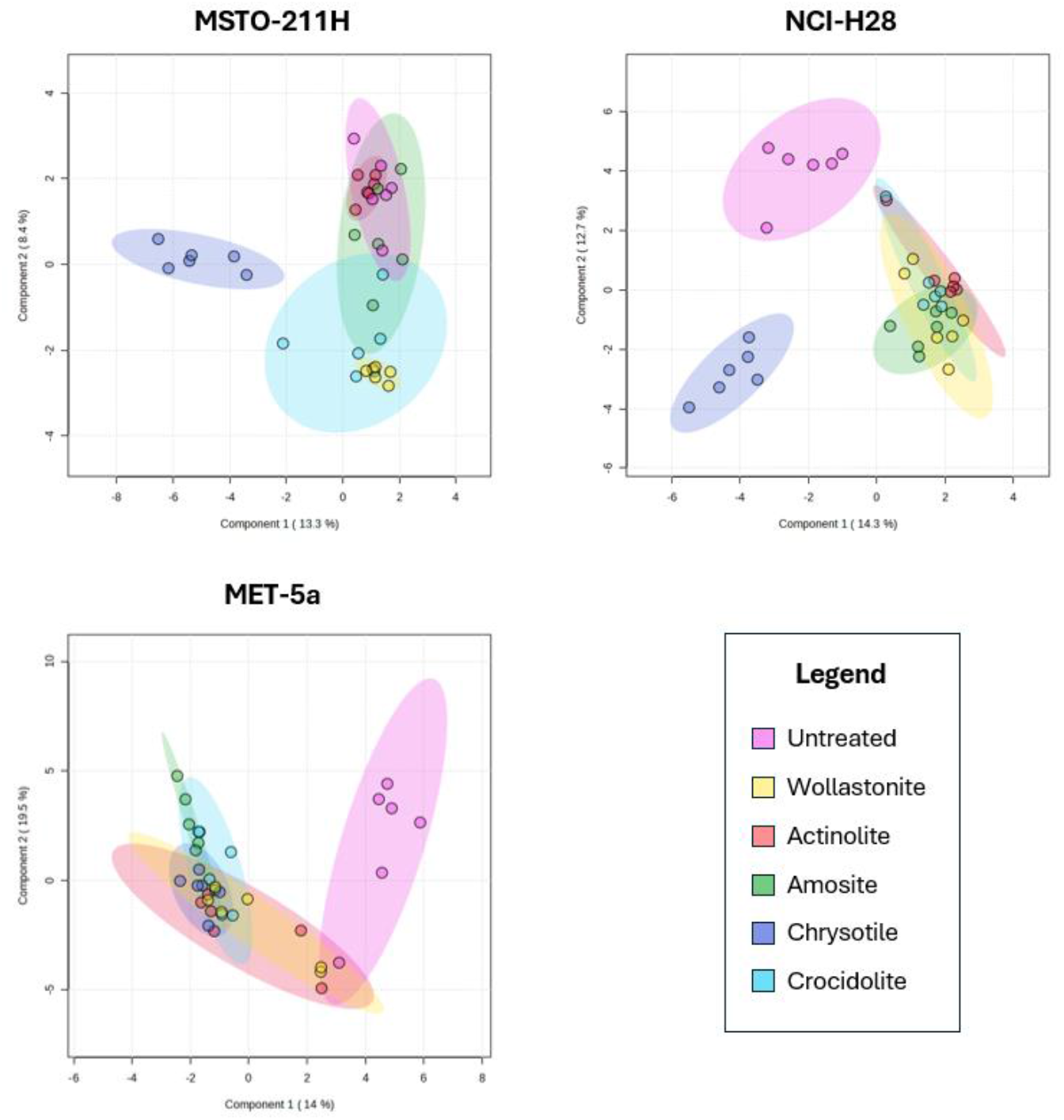
SPLS-DA plots of mesothelial cell lines treated with AMFs. Each individual point represents a single experimental repeat (n=6 per treatment).

The MSTO-211H cell line separated into two groups, one containing only chrysotile treated cells, and the second group consisting of all other AMF treated cells and untreated cells. All treatments in the second group were closely clustered, which suggests that the only AMF treatment which affected the MSTO-211H cell line VOC profile was amosite, as all other treatments resulted in similar grouping following PCA.

Applying sPLS-DA resulted in much greater separation between the groups and a clearer differentiation between untreated and AMF treated VOC profiles across all cell lines (Figure 3).

sPLS-DA resulted in similar clustering and numbers of treatment clusters to PCA. The largest observed difference was the separation of the NCI-H28 AMF treatments into three distinct groups rather than two, with untreated and chrysotile treated cells clearly separating, rather than clustering together as they did following PCA.

### VOCs associated with AMF exposure

Following filtering and normalization of GC-MS results, statistical analysis revealed forty-seven VOCs which were significantly increased or decreased for at least two mineral fiber treatment for the MET-5a cell line. Subsequent filtering of compounds with low quality matches to the MoNA database produced a final panel of 29 VOCs which were significantly altered (P < 0.05 vs untreated) in the MET-5a cell line following AMF stimulation (Table *2*).

The same procedure was followed for the MSTO-211H and NCI-H28 cell lines, where thirteen and twenty-nine significant VOCs were identified respectively (Tables 3 & 4). Individual identities of VOCs varied between each cell line, however significantly altered levels of alkanes, methylated alkanes, alcohols, and aromatic compounds were observed in every cell line following AMF exposure.

**Table 3.**
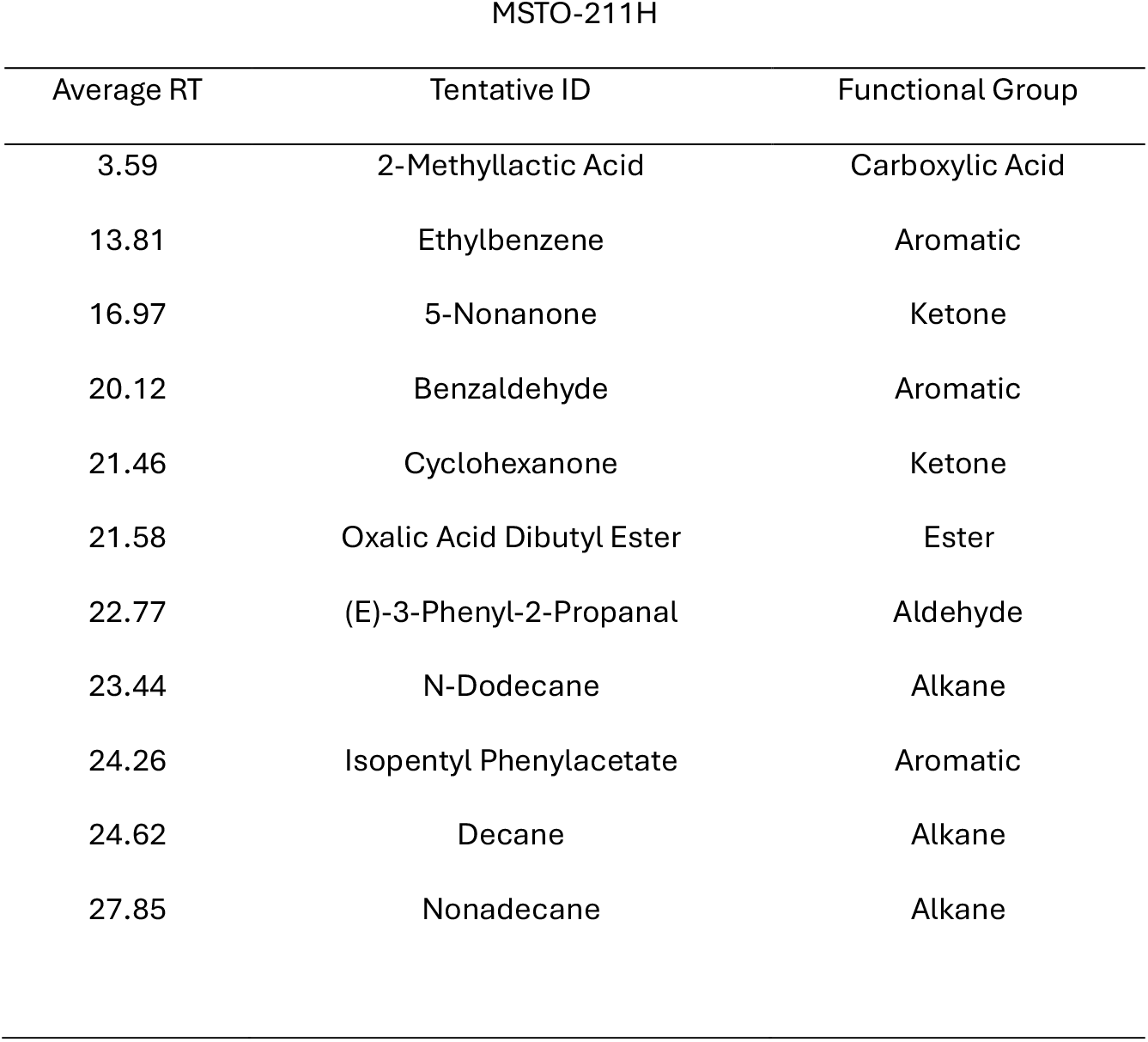
Significantly altered VOCs (P<0.05 vs untreated) in the MSTO-211H cell line following AMF exposure.

**Table 4.** Significantly altered VOCs (P<0.05 vs untreated) in the NCI-H28 cell line following AMF exposure.

| Average RT | Tentative ID | Functional Group |
| --- | --- | --- |
| 1.35 | Dimethylamine | Amine |
| 4.02 | Hexane | Alkane |
| 8.26 | Benzene | Aromatic |
| 13.81 | Isopentyl Phenylacetate | Aromatic |
| 15.76 | Undecane | Alkane |
| 16.19 | 2-Hexanone | Ketone |
| 17.46 | Meta Xylene | Aromatic |
| 18.66 | Butyl Lactate | Ester |
| 19.47 | 1-Hexanol | Alcohol |
| 19.77 | Isopropylbenzene | aromatic |
| 20.11 | 5-Nonanone | Alkane |
| 20.88 | 2-Methylheptane | Alkane |
| 20.88 | Hexamethylethane | Alkane |
| 21.10 | Propylbenzene | Aromatic |
| 21.46 | Decane | Alkane |
| 21.58 | Oxalic Acid Dibutyl Ester | Ester |
| 23.44 | Tridecane | Alkane |
| 23.44 | N-Octadecane | Alkane |
| 24.26 | Benzaldehyde | Aldehyde |
| 25.05 | 2-Allyloxyethanol | Alcohol |
| 25.06 | Methyl Hexyl Ketone | Ketone |
| 25.43 | Isobutylbenzene | Aromatic |
| 25.56 | Hexadecane | Alkane |
| 27.91 | Nonadecane | Alkane |
| 27.99 | Phenol | Aromatic |
| 28.51 | Docosane | Alkane |
| 28.97 | Isobutyl Phenyl Ketone | Ketone |
| 30.80 | 1,2,4-Trichlorobenzene | Aromatic |
| 38.94 | Hexacosane | Alkane |

In summary, a total of four VOCs were identified which were consistently significantly altered following AMF exposure in every cell line (Table 5), however, it should be noted that no VOCs were found which were significantly altered by every AMF treatment in every cell line.

**Table 5.** Significantly altered VOCs (P<0.05 vs untreated) which were detected in every cell line for at least two AMF treatments.

| All Cell Lines |  |
| --- | --- |
| Average RT | Tentative ID |
| 13.81 | Isopentyl Phenylacetate |
| 20.12 | 5-Nonanone |
| 21.34 | Decane |
| 24.25 | Benzaldehyde |

## 4. Discussion

Enhancing the clinical diagnostic options for malignant mesothelioma (MM) remains an urgent priority to improve patient outcomes. The identification of clinically relevant early detection biomarkers for MM would eliminate the need for current invasive procedures such as thoracentesis and pleural biopsy. Here, we have successfully demonstrated VOC metabolites specific to asbestos fiber exposure in a controlled environment, generating potential biomarkers of MM. MM is classified as an inflammatory cancer, and its extended latency period suggests that patients undergo a prolonged period of oxidative stress in the pulmonary region, which may be reflected in an individual’s breath which could be tested for the VOCs presented in this research.

Previous studies [12,13] investigated the VOC profile of MM cell lines *in vitro*. Little *et al* showed that mesothelial cell subtype can be differentiated based on VOC profile, with different mesothelial subtypes, biphasic and epithelioid, having distinct VOC profiles. While the differentiation of mesothelial subtype based on VOC profile represents a step forward in the pursuit of a diagnostic breath test for MM, it is important to recognise the pathogenesis of MM involves a lengthy latency period following initial exposure to asbestos [21]. An ideal breath test for MM would be capable of detecting MM at an earlier, more treatable stage than current diagnostic methods. Previous studies have focused on the differentiation of healthy volunteers from MM patients [8,9,22], however the prolonged period of inflammation arising from asbestos exposure has not been explored in relation to its effect on patient breath VOC profile. Additionally, the effect of AMF exposure on the VOC profile of mammalian cell lines has not been explored either.

By building a VOC profile which can be attributed to AMF exposure, it may be possible to detect MM in asbestos exposed individuals during the latency period. Following asbestos exposure, it is likely that that there is a transitional period as mesothelial cells progress from a healthy state to a malignant state as genetic mutations accumulate [23], which may be reflected in exogenous VOCs arising from an altered metabolic state because of *in situ* AMFs.

This study investigated the impact of four AMFs and one non-AMF on the VOC profiles of mesothelial cell lines. Every mineral fibre treatment was found to induce alterations in VOC profile in every cell line. Despite this, there were differences observed between each AMF treatment in terms of the specific VOCs which were altered. For example, of the twelve total VOCs which were altered in the MSTO-211H cell line, only four VOCs were altered following all AMF treatments, suggesting that these VOCs could function as generalised markers of AMF exposure regardless of the identity of the AMF. Conversely, actinolite and chrysotile AMF exposure induced changes in VOCs which were unique to that AMF treatment. Amosite and chrysotile AMF exposure induced changes in a total of 11 VOCs, whereas the wollastonite fibre treatment induced changes in seventeen VOCs, the most of any AMF treatment.

AMF exposure did not significantly affect live cell counts in these experiments **(Figure 1)**. Variations in the number of viable cells would change the levels and identities of VOCs which are detected [24], the lack of any significant alteration in cellular viability or overall cell count suggests that any alterations in VOC profile following asbestos fibre stimulation are a direct result of asbestos fibre stimulation and not reduced cellular viability.

The initial statistical analysis of significantly altered compounds revealed a panel of forty one unique VOCs associated with AMF exposure across the three cell lines. Most of the identified VOCs were long chain alkanes, aromatics and alcohols, classes of compounds which have been previously linked to lipid peroxidation and oxidative stress [25]

Interestingly, wollastonite, the non-AMF control, still induced significant alterations in cell line VOC profiles, resulting in overall VOC profiles like those produced by asbestos fibre stimulation. It is plausible that the physical presence of mineral fibres within and in proximity to cells is responsible for the metabolic response which induces an altered

VOC profile, rather than a specific molecular or structural property of asbestos itself. While wollastonite is not categorized as an asbestiform fibre, it induces similar changes in cellular VOC profile to AMFs as shown by PCA and SPLS-DA plots (Figures 2 & 3).

Previous small-scale studies have successfully differentiated healthy volunteers from asbestos exposed (Aex) individuals and MM patients based on breath VOC profile [26-29], which supports the notion that *in situ* AMFs within the mesothelium may induce, or contribute to, a detectable change in breath VOC profile long before malignancy develops.

Four VOCs were found to be significantly altered in every cell line following AMF stimulation (Table 5), indicating that these specific compounds could potentially serve as general markers for asbestos exposure.

Classification of the identified VOCs into their functional groups aids the understanding of how AMFs influence cellular VOC profile. Many studies cite the identities of individual VOCs as individual biomarkers for mesothelioma; however, it is unlikely that a single VOC can function as a lone biomarker for disease. Rather, an overall profile attributed to specific classes of VOCs may be more clinically relevant [25]. Human metabolism is complex, and metabolic pathways involving lipid peroxidation arising from oxidative stress can result in a multitude of VOC products which can be linked back to the initial oxidative stress inducing factor. We have identified alkanes, methylated alkanes, alcohols, ketones and aromatics as five classes of exogenous VOCs which are significantly affected by AMF exposure (Tables 2, 3 & 4). Future studies may include the deeper investigation of the exact mechanisms responsible for their alteration via asbestos exposure.

The trends seen in volatile levels in mesothelial cell lines following AMF exposure point towards the alteration of endogenous levels of specific classes of VOCs, rather than an increase or decrease in a single VOC. Every AMF treatment induced changes in at least one VOC belonging to these classes of compounds in every cell line. While the specific identity of VOCs varied between each cell line, the impact on VOC classes was similar for all cell lines. For example, it is unlikely that there is a major difference between the release of pentadecane and tetradecane metabolically, as they are both long chain alkanes differing by the incorporation of a single carbon. It is more likely that the presence of long chain alkanes in general is indicative of oxidative stress [25] because of acute AMF exposure.

While this study represents an important first step in understanding the relationship between AMF exposure and cellular VOC profiles, it is important to acknowledge some limitations. While the MET-5a cell line used in this study is classified as non-malignant, this cell line is rendered immortal via SV40 transformation, resulting in total deletion of p53. Such alterations to a fundamental tumour suppressor gene should be considered when classifying this cell line as a model of normal mesothelial cells (ATCC, CRL-9444).

Comparison of the effects of AMF stimulation on non-malignant and malignant cell lines is important as it shows how aberrant genetic states may impact the VOCs released because of oxidative stress. AMFs are biopersistent and remain *in situ* in the mesothelium [30], therefore it is likely that MM patient breath VOC profiles will continue to be affected by AMF stimulation following transformation to a malignant state. While chrysotile has been shown to not have the same biopersistence as amphibole AMFs [31], exposure to AMFs in general may still result in an altered VOC profile. The MET-5a cell line is a non-malignant, immortalized cell line which can be used as a model for normal mesothelial cells *in vitro*. The effects of AMF stimulation on MET-5a cell lines may be analogous to the effect of AMF fibres on healthy human mesothelial tissue.

## 5. Conclusions

This study exposed cell lines to AMFs for a period of 48 hours. Compared to the latency period of MM, this is a short period of time, however this was sufficient to induce detectable changes to cellular VOC profile. Additionally, previous studies investigating the effects of AMF and AMF-like fibres on mesothelial cell lines showed that an exposure time of 48 hours was sufficient to induce detectable transcriptomic changes [32,33]. Longer term AMF exposure to mesothelial cell lines would be a logical next step and would reveal how cellular VOC profile may change in response to longer term exposure.

In conclusion, we have shown that AMF stimulation significantly affects the VOC profile of both malignant and non-malignant mesothelial cell lines. Data identifies and supports a panel of VOCs which correspond to general AMF exposure. Our results agree with a recent systematic review which identified classes of compounds that are significantly altered in a state of oxidative stress are primarily long chain alkanes, aromatic compounds, and alcohols [25]. The results presented herein may be applicable to human volunteer studies, as these VOCs may also be indicators of asbestos exposure in asbestos exposed individuals.

## Data availability statement

The datasets generated during and/or analysed during the current study are available from the corresponding author on reasonable request.

## Acknowledgements

This research is supported by a UKRI PhD Studentship from Sheffield Hallam University (SB, SH-S) and a Cancer Research UK Early Detection and Diagnosis Primer Award, reference EDDPMA-May22\100084 (TI, SH-S).

## Author Contribution

Conceptualization, S.HS, N.P, J.W, and M.H.; Data curation, S.B.; Formal analysis, S.B, T.I.; Funding acquisition, S.HS., N.P., J.W., and M.H.; Investigation, S.B.; Methodology, S.B and S.HS.; Project administration, S.HS., J.W., M.H and N.P.; Resources, W.J.B.; Visualization, S.B.; Writing—original draft, S.B.; Writing—review and editing, S.B., T.I., S.HS, N.P, J.W, and M.H. All authors have read and agreed to the published version of the manuscript.

## Conflict of interest statemen

All authors declare that there are no conflicts of interest.

